# Improving on a modal-based estimation method: model averaging for consistent and efficient estimation in Mendelian randomization when a plurality of candidate instruments are valid

**DOI:** 10.1101/175372

**Authors:** Stephen Burgess, Verena Zuber, Apostolos Gkatzionis, Jessica MB Rees, Christopher N Foley

## Abstract

**Background:** A robust method for Mendelian randomization does not require all genetic variants to be valid instruments to give consistent estimates of a causal parameter. Several such methods have been developed, including a mode-based estimation method giving consistent estimates if a plurality of genetic variants are valid instruments; that is, there is no larger subset of invalid instruments estimating the same causal parameter than the subset of valid instruments.

**Methods:** We here develop a model averaging method that gives consistent estimates under the same ‘plurality of valid instruments’ assumption. The method considers a mixture distribution of estimates derived from each subset of genetic variants. The estimates are weighted such that subsets with more genetic variants receive more weight, unless variants in the subset have heterogeneous causal estimates, in which case that subset is severely downweighted. The mode of this mixture distribution is the causal estimate. This heterogeneity-penalized model averaging method has several technical advantages over the previously proposed mode-based estimation method.

**Results:** The heterogeneity-penalized model averaging method outperformed the mode-based estimation in terms of efficiency and outperformed other robust methods in terms of Type 1 error rate in an extensive simulation analysis. The proposed method suggests two distinct mechanisms by which inflammation affects coronary heart disease risk, with subsets of variants suggesting both positive and negative causal effects.

**Conclusions:** The heterogeneity-penalized model averaging method is an additional robust method for Mendelian randomization with excellent theoretical and practical properties, and can reveal features in the data such as the presence of multiple causal mechanisms. (249 words)

**Key messages:**

- We propose a heterogeneity-penalized model averaging method that gives consistent causal estimates if a weighted plurality of the genetic variants are valid instruments.
- The method calculates causal estimates based on all subsets of genetic variants, and upweights subsets containing several genetic variants with similar causal estimates.
- The method is asymptotically efficient and does not rely on bootstrapping to obtain a confidence interval, nor is the confidence interval constrained to be symmetric.
- In particular, the confidence interval can include multiple disjoint intervals, suggesting the presence of multiple causal mechanisms by which the risk factor influences the outcome.
- The method can incorporate biological knowledge to upweight the contribution of genetic variants with stronger plausibility of being valid instruments.

## Introduction

Mendelian randomization is an epidemiological approach for making causal inferences from observational data by using genetic variants as instrumental variables [1, 2]. If a genetic variant is a valid instrument for the risk factor, then any association of the variant with the outcome is indicative of a causal effect of the risk factor on the outcome [3]. When there are multiple genetic variants that are all valid instrumental variables, and under certain parametric assumptions (most notably that all relationships between variables are linear and there is no effect modification), an efficient test of the causal null hypothesis as the sample size increases can be obtained using the two-stage least squares method (based on individual-level data) [4] or equivalently the inverse-variance weighted (IVW) method (based on summarized data) [5]. With uncorrelated instruments, the IVW estimate (equal to the two-stage least squares (2SLS) estimate) is a weighted mean of the Wald (or ratio) estimates obtained separately from each individual instrumental variable.

While the 2SLS/IVW estimator is asymptotically efficient, it is not robust to violations of the instrumental variable assumptions. Specifically, if a genetic variant is a valid instrument, then the ratio estimate based on that variant is a consistent estimate of the causal effect. Hence the weighted mean of these ratio estimates is a consistent estimate of the causal effect if all genetic variants are valid instruments, but not in general if at least one variant is not a valid instrument [6]. This has motivated the development of robust methods for instrumental variable analysis based on only a subset of the genetic variants being valid instruments. For example, Kang et al. developed a method using L1-penalization that gives consistent estimates if at least 50% of the instrumental variables are valid [7]. Bowden et al. considered simple and weighted median methods that again are consistent if at least 50% of the instrumental variables are valid; the simple median method is a median of the variant-specific ratio estimates [8]. Most recently, Hartwig et al. have developed a modal-based estimation method that provides a consistent estimate under the zero modal pleiotropy assumption (ZEMPA) [9]. This assumption states that, in large sample sizes, the largest subset of variants with the same ratio estimate comprises the valid instruments. Invalid instruments may have different ratio estimates asymptotically, but there is no larger subset of invalid instruments with the same ratio estimate than the subset of valid instruments. Intuitively, this means that the true causal estimate can be identified asymptotically as the mode of the variant-specific ratio estimates.

While the idea of a modal-based estimate has merit, there are several issues with the implementation of Hartwig’s modal-based estimate that could be improved upon. In particular, their implementation of this approach fits a kernel density-smoothed function to the variant-specific ratio estimates, and calculates confidence intervals based on the median absolute deviation of a bootstrapped distribution. Varying the bandwidth of the kernel density can result in substantial changes to the estimate and its confidence interval, as demonstrated later in this paper.

In this paper, we propose an alternative way of constructing a density function for the causal effect estimate as a heterogeneity-penalized weighted mixture distribution. This approach upweights estimates that are supported by multiple genetic variants, but severely downweights heterogeneity. We show that the mode of this distribution will be an asymptotically consistent estimator of the causal effect if a weighted plurality of the genetic variants are valid instruments. We first introduce this method, and then we demonstrate its performance in a simulation study compared to other robust methods. We consider its behaviour in two applied examples. Finally, we discuss the results of this paper and their relevance to applied research. In particular, we consider how to incorporate biological knowledge into the weighting procedure. Software code for implementing the proposed method is provided in the Supplementary Material.

## Methods

In this section, we first introduce the data requirements and parametric assumptions necessary for summarized data Mendelian randomization. We then recall the inverse-variance weighted method, and subsequently introduce the model averaging procedure proposed in this paper.

### Data requirements and assumptions

For practical reasons, many modern Mendelian randomization investigations are conducted using summarized data on genetic associations with the risk factor (*X*) and outcome (*Y*) taken from univariable regression models of the risk factor (or outcome) regressed on the genetic variants in turn [10]. We assume, as is common in applied practice, that the genetic variants are all uncorrelated (not in linkage disequilibrium). For each genetic variant *G*_*j*_ (*j* = 1, 2*,…, J*), we assume that we have an estimate 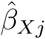 of the association of the genetic variant with the risk factor obtained from linear regression. Similar association estimates are assumed to be available for the outcome 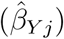. The standard error of the association estimate with the outcome is 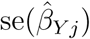. If any of the variables is binary, then these summarized association estimates may be replaced with association estimates from logistic regression; as has been shown previously, the interpretation of the causal estimate in this case is not clear due to non-collapsibility, but estimates still represent valid tests of the causal null hypothesis [11, 12]. See Bowden et al. [13] for a more detailed exposition of the parametric assumptions typically made in summarized data Mendelian randomization investigations that are also made here.

### Inverse-variance weighted method

The ratio estimate based on genetic variant *j* is 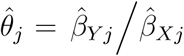, with standard error taken as 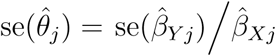 (the leading order term from the delta expansion for the standard error of the ratio of two variables). The inverse-variance weighted (IVW) estimate is a weighted mean of the ratio estimates:

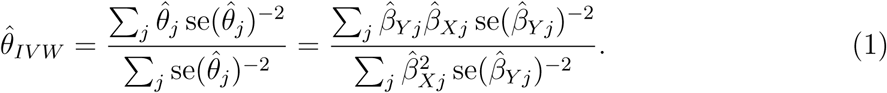

The same estimate can be obtained from the weighted regression:

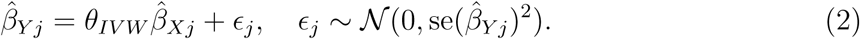

For uncorrelated variants, this estimate is also equivalent to the estimate obtained from two-stage least squares, a method typically used for instrumental variable analysis with individual-level data [5]. These estimates do not take into account uncertainty in the genetic associations with the risk factor; however, these associations are typically more precisely estimated than those with the outcome, and ignoring this uncertainty does not lead to inflated Type 1 error rates in realistic scenarios [14].

The standard error of the IVW estimate based on a fixed-effect meta-analysis model is:

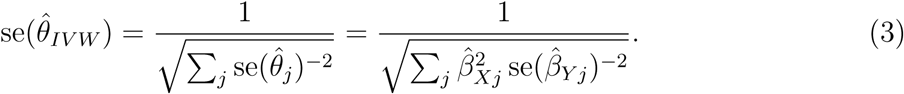

We also consider a multiplicative random-effects model based on the weighted linear regression above:

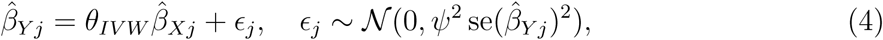

where *ψ* is the residual standard error. Most statistical software packages estimate this additional parameter by default in a weighted linear regression model. A fixed-effect analysis can be performed by fixing the value of *ψ* to 1 [15]. To ensure that the standard error of the IVW estimate is never more precise than that from a fixed-effect analysis, we allow *ψ* to take values above 1 (corresponding to over-dispersion of the genetic association estimates), but not values below 1 (corresponding to under-dispersion). If all genetic variants estimate the same causal parameter, then *ψ* should tend to 1 asymptotically.

### Heterogeneity-penalized model averaging method

We seek to estimate a distribution with the property that the mode (the maximum value) of the distribution will tend to the true causal effect when a plurality of the genetic variants are valid instruments. We consider a model averaging procedure with 2^*J*^ − *J* − 1 candidate models, where *J* is the total number of genetic variants. Each model corresponds to one of the 2^*J*^ − J − 1 subsets of genetic variants (subsets including 0 or 1 genetic variants are ignored throughout). We consider a mixture distribution of 2^*J*^ − J − 1 normal distributions, where the *k*th normal distribution has mean and standard deviation corresponding to the IVW estimate and standard error based on all the variants in the *k*th subset:

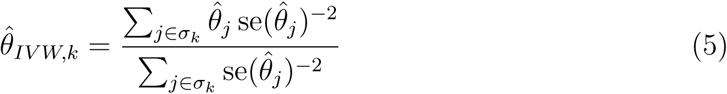

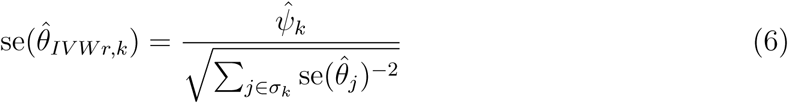

where *σ*_*k*_ = (*σ*_*k*1_*, σ*_*k*2_*,…, σ*_*kJ*_) : *σ*_*kj*_ *∈* {0, 1} represents a subset of the genetic variants, *j ∈ σ*_*k*_ when *σ*_*kj*_ = 1 (this means that 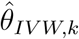 is the IVW estimate based on all the variants in subset *k*), and

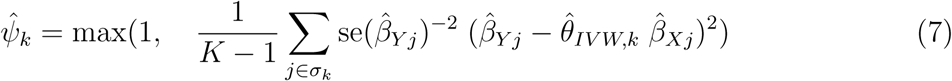

where *K* is the number of variants included in subset *k*. The random-effects versions of the standard error 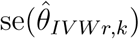 are used in this mixture distribution to appropriately allow for heterogeneity between the variant-specific ratio estimates in the overall causal estimate.

The weight given to each of these normal distributions is calculated as:

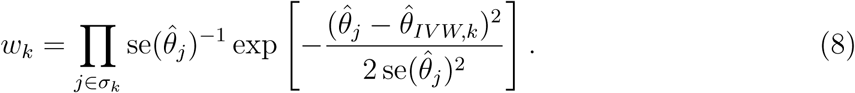

Aside from the constant term, this is the likelihood assuming the variant-specific ratio estimates 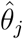 are normally distributed about a common mean 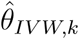 with variant-specific standard deviation 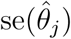. Weights will be larger when more variants are included in the subset *k* due to the 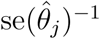 terms, but they will reduce sharply if there is more heterogeneity between the variant-specific ratio estimates for variants in the subset than would be expected due to statistical uncertainty alone if all variants estimated the same causal parameter. If the variant-specific ratio estimates for variants in a particular subset substantially differ, then the weight for that subset will be low. Note that the reason for excluding subsets with one variant is that heterogeneity cannot be estimated for these subsets. We then normalize the weights so that they sum to one:

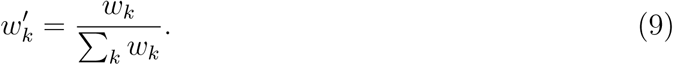

The causal estimate is the mode of the the mixture of normal distributions using these weights:

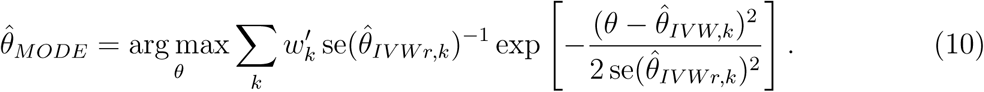

### Consistency and efficiency

In the asymptotic limit for a fixed number of genetic variants but as the sample size tends to infinity (and hence the standard errors of the ratio estimates decrease to zero), the weighted mixture distribution tends to a series of spikes about the IVW estimates based on each subset of variants. The height of each spike depends on the total weight of variants that have that causal estimate, and the tallest spike is the estimate with the greatest weight of evidence. The modal estimate will be the IVW estimate corresponding to the subset *k* of variants all having the same ratio estimate which has the greatest product of the inverse standard errors of the ratio estimates 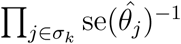. Therefore a consistent estimate is obtained under a Hartwig’s weighted ZEMPA assumption [9]. The intuition of this assumption is that a weighted plurality of the genetic variants is required to be valid instruments (as opposed to median-based methods that require a majority or weighted majority of variants to be valid instruments). The term ‘plurality’ is taken from the terminology of elections; a political party winning more votes than any other is said to have a plurality of the votes.

Under this assumption, the heterogeneity-penalized model averaging method is asymptotically efficient, as the weight of the IVW estimate based on all the valid instruments will increase to 1 as the sample size tends to infinity. This can be seen as the weight for any subset containing variants with different ratio estimates will decrease to zero rapidly. The weight of the largest subset of variants with the same ratio estimates will be the greatest of all subsets by the ZEMPA assumption, and the ratio of this weight to all other weights will increase to infinity as the sample size increases. However, asymptotic efficiency is not necessarily an important property in practice, as infinite sample sizes are rarely encountered in applied investigations. The model averaging estimate should be efficient for finite sample sizes when several variants have similar ratio estimates.

### Inferences on the weighted model-averaged distribution

We perform causal inferences based on the model-averaged distribution using a generalized likelihood ratio test to construct a confidence interval. We take twice the log-likelihood function, and construct a confidence interval consisting of all points for which twice their log-likelihood is within a given vertical distance from the modal estimate. For a 95% confidence interval, this distance is 3.841 (half of the 95th percentile of a chi-squared distribution with 1 degree of freedom). This is based on the result that twice the difference in the log-likelihood at the estimate and at the true value of the parameter has a chi-squared distribution (here with 1 degree of freedom as the parameter is 1-dimensional). This results in inference without requiring resampling techniques (such as bootstrapping). The confidence interval is not guaranteed to be symmetrical, or to be a single range of values (see later for an example of a bimodal weighted distribution resulting in a composite confidence interval).

Practically, the modal estimate and confidence interval were obtained using a grid search approach. The likelihood was evaluated at a series of points (in the simulation study, from −1 to +1 at intervals of 0.001 – so estimates and confidence intervals were estimated to 3 decimal places). The modal estimate was taken as the point with the greatest value of the likelihood function, and the 95% confidence interval was taken as the set of points for which twice the log-likelihood was within 3.841 of the twice the log-likelihood at the modal estimate.

## Simulation study

To consider the expected performance of this proposed method in realistic situations as well as in comparison to alternative robust methods, we perform a simulation study. We consider four scenarios:

1. no pleiotropy – all genetic variants are valid instruments;
2. balanced pleiotropy – some genetic variants have direct (pleiotropic) effects on the outcome, and these pleiotropic effects are equally likely to be positive as negative;
3. directional pleiotropy – some genetic variants have direct (pleiotropic) effects on the outcome, and these pleiotropic effects are simulated to be positive;
4. directional pleiotropy via a confounder – some genetic variants have pleiotropic effects on the outcome via a confounder. These pleiotropic effects are correlated with the instrument strength.

In the first three scenarios, the Instrument Strength Independent of Direct Effect (InSIDE) assumption [6] is satisfied; in Scenario 4, it is violated. This is the assumption required for the MR-Egger method to provide consistent estimates.

We simulate data for a risk factor *X*, outcome *Y*, confounder *U* (assumed unmeasured), and *J* genetic variants *G*_*j*_*, j* = 1*,…, J*. Individuals are indexed by *i*. The data-generating model for the simulation study is as follows:

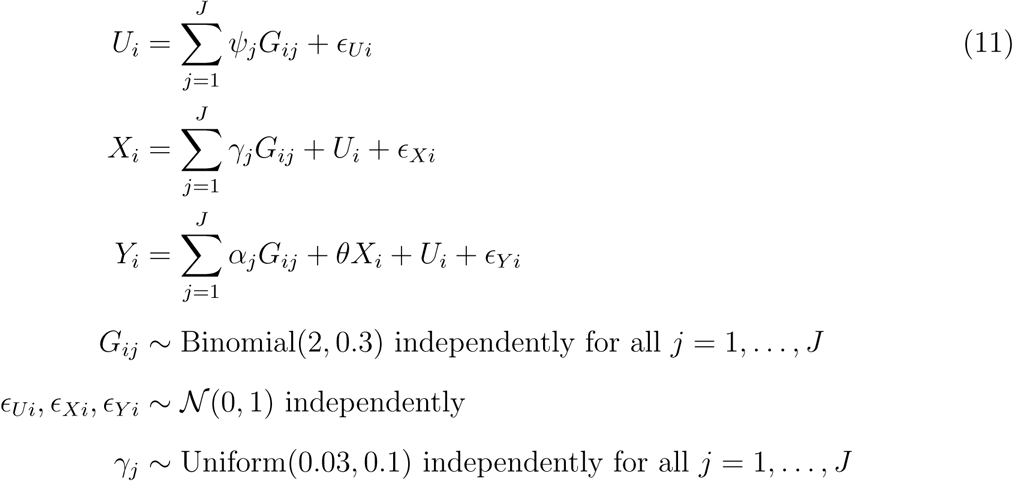

The risk factor and outcome are positively correlated due to confounding even when the causal effect *θ* is zero through the unmeasured confounder *U*. The genetic variants are modelled as single nucleotide polymorphisms (SNPs) with a minor allele frequency of 30%. A total of *J* = 10 genetic variants are used in each analysis. As the proposed model-averaging method calculates weights for all 2^*J*^ *- J -* 1 possible models, the model scales exponentially with the number of variants, and so including more variants was not computationally feasible in a simulation setting. For each of Scenarios 2 to 4, we considered cases with 2, 3 and 5 invalid instruments. For valid instruments, the *α*_*j*_ and *ψ*_*j*_ parameters were set to zero. For invalid instruments, the *α*_*j*_ parameters were either drawn from a uniform distribution on the interval from *-*0.1 to 0.1 (Scenario 2), or from 0 to 0.1 (Scenario 3), or set to zero (Scenario 4). The *ψ*_*j*_ parameters were either set to zero (Scenarios 2 and 3), or drawn from a uniform distribution on the interval from *-*0.1 to 0.1 (Scenario 4). The causal effect *θ* was either set to 0 (no causal effect) or 0.2 (positive causal effect). The average value of the *R*^2^ statistic for the 10 variants across simulated datasets was 1.0% (from 1.1 to 1.4% in Scenario 4) corresponding to an average F statistic of 20.4 (from 23.4 to 27.5 in Scenario 4).

In total, 10 000 datasets were generated in each scenario. We considered a two-sample setting in which genetic associations with the risk factor and outcome were estimated on non-overlapping groups of 20 000 individuals. We compared estimates from the proposed heterogeneity-penalized model averaging method with those from a variety of methods: the standard IVW method, MR-Egger [6] (both using random-effects), weighted and simple median [8], and the mode-based estimate (MBE) of Hartwig et al. [9]. Each of the methods was implemented using summarized data only.

### Results

Results for all of the methods are provided in Tables 1 (Scenario 1) and 2 (Scenarios 2 to 4). We provide the mean estimate, the standard deviation of estimates, the mean standard error (Table 1 only), and the empirical power of the 95% confidence interval (the proportion of 95% confidence intervals excluding the null; this is the Type 1 error rate with a null causal effect). Results for the MBE method are only provided for 1000 simulated datasets per scenario. This is for computational reasons – the MBE method took around 20 times longer to run than all the other methods put together. Results for the MBE method correspond to simple (unweighted) and weighted versions of the method not assuming NOME (no measurement error) with the recommended bandwidth parameter from the modified Silverman rule (*ϕ* = 1) [16]; in total, 12 different versions of the MBE method are proposed by Hartwig et al.

**Table 1.** Mean, standard deviation (SD), mean standard error (mean SE) of estimates, and empirical power (%) for Scenario 1 (all variants valid instruments).

| Method | Scenario 1: all instruments valid |  |  |  |
| --- | --- | --- | --- | --- |
|  | Mean | SD | Mean SE | Power |
| Null causal effect: $\theta = 0$ | | | | |
| Inverse-variance weighted | 0.001 | 0.072 | 0.077 | 3.9 |
| MR-Egger | 0.003 | 0.223 | 0.236 | 3.6 |
| Simple median | 0.001 | 0.092 | 0.105 | 2.1 |
| Weighted median | 0.002 | 0.086 | 0.096 | 2.8 |
| Simple mode-based estimate (Hartwig) | 0.003 | 0.113 | 0.149 | 0.3 |
| Weighted mode-based estimate (Hartwig) | 0.002 | 0.098 | 0.128 | 1.2 |
| Heterogeneity-penalized model averaging | 0.001 | 0.080 | - | 1.4 |
| Positive causal effect: $\theta = +0.2$ | | | | |
| Inverse-variance weighted | 0.191 | 0.080 | 0.086 | 61.9 |
| MR-Egger | 0.130 | 0.250 | 0.263 | 7.0 |
| Simple median | 0.201 | 0.104 | 0.119 | 39.0 |
| Weighted median | 0.185 | 0.096 | 0.109 | 39.9 |
| Simple mode-based estimate (Hartwig) | 0.195 | 0.136 | 0.167 | 18.5 |
| Weighted mode-based estimate (Hartwig) | 0.172 | 0.115 | 0.142 | 22.4 |
| Heterogeneity-penalized model averaging | 0.188 | 0.090 | - | 38.8 |

Table 1 shows the efficiency of the model averaging method when all genetic variants are valid instruments. The method is considerably more efficient than the MR-Egger and MBE methods, with less variable estimates and greater power to detect a causal effect, and similar in efficiency to the median-based methods. Coverage under the null is conservative for all methods, but particularly for the MBE and model averaging methods.

Table 2 shows the robustness of the model averaging method in a range of invalid instrument scenarios. Type 1 error rates are well-controlled (less than 7.5%) in all scenarios when 2 or 3 out of the 10 variants are invalid, and generally below those of other methods even when 5 variants are invalid. Compared with the model averaging method, Type 1 error rates with 5 invalid instruments for the MR-Egger method are lower in Scenario 3; however, they are far higher in Scenario 4, and the power of the MR-Egger method to detect a positive causal effect was low throughout. Equally, Type 1 error rates are slightly lower for the simple median method in Scenario 4, but higher in Scenario 3. The empirical power of the model averaging method to detect a causal effect was generally lower than that for other methods. However, when a method suffers from Type 1 error inflation, this comparison is not a fair one. The power of the model averaging method to detect a positive causal effect was not dominated by any method that had well-controlled Type 1 error rates. Indeed, in Scenario 2, the power of the model averaging method even exceeded that of the IVW method with 3 and 5 invalid variants. This is because models including the invalid variants are downweighted in the model averaging method, whereas these variants inflate the standard error in the IVW method.

**Table 2.** Mean, standard deviation (SD) of estimates, and empirical power (%) for scenarios 2, 3, and 4. Abbreviation: MBE = mode-based estimate of Hartwig et al. [9].

| Method | 2 invalid variants |  |  | 3 invalid variants |  |  | 5 invalid variants |  |  |
| --- | --- | --- | --- | --- | --- | --- | --- | --- | --- |
|  | Mean | SD | Power | Mean | SD | Power | Mean | SD | Power |
| Null causal effect: $\theta = 0$ | | | | | | | | | |
| Scenario 2: Balanced pleiotropy, InSIDE satisfied |  |  |  |  |  |  |  |  |  |
| Inverse-variance weighted | -0.001 | 0.140 | 6.3 | 0.002 | 0.163 | 7.5 | 0.000 | 0.202 | 7.8 |
| MR-Egger | 0.001 | 0.436 | 7.7 | 0.004 | 0.509 | 8.2 | 0.007 | 0.629 | 9.3 |
| Simple median | 0.000 | 0.113 | 3.8 | 0.002 | 0.129 | 5.5 | 0.000 | 0.175 | 10.2 |
| Weighted median | 0.001 | 0.109 | 5.2 | 0.001 | 0.125 | 7.5 | 0.000 | 0.178 | 15.0 |
| Simple MBE | 0.000 | 0.126 | 1.0 | 0.008 | 0.131 | 1.8 | 0.006 | 0.196 | 4.0 |
| Weighted MBE | 0.004 | 0.105 | 2.4 | 0.000 | 0.113 | 3.1 | 0.005 | 0.172 | 8.3 |
| Model averaging | 0.000 | 0.100 | 2.4 | 0.000 | 0.115 | 3.2 | -0.001 | 0.187 | 6.0 |
| Scenario 3: Directional pleiotropy, InSIDE satisfied |  |  |  |  |  |  |  |  |  |
| Inverse-variance weighted | 0.136 | 0.101 | 10.8 | 0.206 | 0.113 | 20.9 | 0.342 | 0.131 | 52.2 |
| MR-Egger | 0.004 | 0.421 | 7.8 | 0.002 | 0.479 | 8.2 | 0.011 | 0.539 | 8.5 |
| Simple median | 0.065 | 0.104 | 5.2 | 0.113 | 0.118 | 11.1 | 0.273 | 0.172 | 44.5 |
| Weighted median | 0.054 | 0.104 | 6.9 | 0.096 | 0.123 | 13.1 | 0.225 | 0.182 | 40.9 |
| Simple MBE | 0.020 | 0.122 | 1.7 | 0.044 | 0.138 | 2.3 | 0.146 | 0.220 | 9.4 |
| Weighted MBE | 0.013 | 0.102 | 2.9 | 0.041 | 0.123 | 5.1 | 0.114 | 0.177 | 12.8 |
| Model averaging | 0.021 | 0.098 | 2.6 | 0.043 | 0.121 | 3.9 | 0.133 | 0.214 | 11.8 |
| Scenario 4: Directional pleiotropy, InSIDE violated |  |  |  |  |  |  |  |  |  |
| Inverse-variance weighted | 0.104 | 0.125 | 19.4 | 0.150 | 0.135 | 26.2 | 0.232 | 0.140 | 38.3 |
| MR-Egger | 0.240 | 0.433 | 35.9 | 0.304 | 0.440 | 39.0 | 0.401 | 0.411 | 40.7 |
| Simple median | 0.023 | 0.111 | 4.1 | 0.044 | 0.125 | 6.5 | 0.095 | 0.164 | 16.9 |
| Weighted median | 0.090 | 0.144 | 20.8 | 0.143 | 0.164 | 34.1 | 0.247 | 0.178 | 60.5 |
| Simple MBE | 0.018 | 0.133 | 2.6 | 0.043 | 0.155 | 4.5 | 0.091 | 0.194 | 12.5 |
| Weighted MBE | 0.072 | 0.171 | 16.4 | 0.128 | 0.197 | 28.2 | 0.216 | 0.204 | 47.6 |
| Model averaging | 0.023 | 0.118 | 4.3 | 0.050 | 0.146 | 7.4 | 0.139 | 0.206 | 22.1 |
| Positive causal effect: $\theta = +0.2$ | | | | | | | | | |
| Scenario 2: Balanced pleiotropy, InSIDE satisfied |  |  |  |  |  |  |  |  |  |
| Inverse-variance weighted | 0.193 | 0.143 | 33.3 | 0.188 | 0.168 | 26.5 | 0.195 | 0.206 | 19.5 |
| MR-Egger | 0.129 | 0.452 | 9.4 | 0.137 | 0.526 | 9.6 | 0.135 | 0.644 | 8.9 |
| Simple median | 0.204 | 0.127 | 34.6 | 0.200 | 0.143 | 33.2 | 0.206 | 0.191 | 33.0 |
| Weighted median | 0.186 | 0.122 | 36.4 | 0.186 | 0.140 | 36.2 | 0.190 | 0.188 | 37.0 |
| Simple MBE | 0.198 | 0.139 | 17.2 | 0.193 | 0.156 | 19.5 | 0.202 | 0.205 | 18.1 |
| Weighted MBE | 0.173 | 0.118 | 21.1 | 0.166 | 0.132 | 22.7 | 0.154 | 0.166 | 21.9 |
| Model averaging | 0.189 | 0.115 | 31.8 | 0.189 | 0.135 | 29.5 | 0.193 | 0.207 | 25.6 |
| Scenario 3: Directional pleiotropy, InSIDE satisfied |  |  |  |  |  |  |  |  |  |
| Inverse-variance weighted | 0.329 | 0.110 | 72.7 | 0.397 | 0.121 | 79.8 | 0.532 | 0.140 | 92.1 |
| MR-Egger | 0.138 | 0.432 | 9.5 | 0.140 | 0.486 | 9.8 | 0.136 | 0.552 | 9.4 |
| Simple median | 0.274 | 0.120 | 55.0 | 0.328 | 0.136 | 65.7 | 0.489 | 0.186 | 87.2 |
| Weighted median | 0.247 | 0.117 | 55.3 | 0.292 | 0.137 | 65.0 | 0.419 | 0.189 | 82.6 |
| Simple MBE | 0.216 | 0.141 | 20.8 | 0.254 | 0.154 | 26.1 | 0.356 | 0.226 | 39.3 |
| Weighted MBE | 0.187 | 0.117 | 24.8 | 0.211 | 0.122 | 31.0 | 0.283 | 0.165 | 48.0 |
| Model averaging | 0.218 | 0.116 | 41.8 | 0.243 | 0.136 | 43.9 | 0.339 | 0.218 | 52.6 |
| Scenario 4: Directional pleiotropy, InSIDE violated |  |  |  |  |  |  |  |  |  |
| Inverse-variance weighted | 0.298 | 0.131 | 63.5 | 0.343 | 0.140 | 66.6 | 0.426 | 0.146 | 74.4 |
| MR-Egger | 0.396 | 0.449 | 42.8 | 0.473 | 0.454 | 48.4 | 0.586 | 0.415 | 51.9 |
| Simple median | 0.232 | 0.125 | 42.7 | 0.252 | 0.139 | 45.7 | 0.304 | 0.176 | 53.2 |
| Weighted median | 0.285 | 0.156 | 62.1 | 0.338 | 0.175 | 71.5 | 0.444 | 0.184 | 85.4 |
| Simple MBE | 0.212 | 0.145 | 22.0 | 0.237 | 0.155 | 25.2 | 0.290 | 0.175 | 37.2 |
| Weighted MBE | 0.245 | 0.173 | 37.1 | 0.293 | 0.195 | 46.8 | 0.383 | 0.202 | 65.4 |
| Model averaging | 0.226 | 0.137 | 40.5 | 0.257 | 0.167 | 42.7 | 0.348 | 0.217 | 52.3 |

In comparison to the MBE method of Hartwig et al., Type 1 error rates for the model averaging method were slightly higher than those for the simple MBE method, but lower than those for the weighted MBE method; particularly in Scenario 4, where the Type 1 error rate for the weighted MBE method was not well-controlled even with only 2 invalid instruments. Power to detect a positive causal effect was greater for the model averaging than for the simple MBE method in all cases, and greater than for the weighted MBE method in all cases except in Scenario 4, where the weighted MBE method had inflated Type 1 error rates. Similar patterns were observed in the bias of estimates, with the model averaging method generally having low bias. Although some methods were less biased in particular scenarios, no method was less biased across all scenarios.

## Applied examples

We provide further illustration of the proposed model averaging method and other robust methods in two applied examples. In the first example, all the variants have similar ratio estimates, whereas in the second example, there is marked heterogeneity in the ratio estimates. Further detail about the applied examples is given in the Supplementary Material.

### Low-density lipoprotein cholesterol and CAD risk

We consider the causal relationship between low-density lipoprotein (LDL) cholesterol and coronary artery disease (CAD) risk based on 8 genetic variants having strong biological links with LDL-cholesterol. Each of these variants is located in a gene region that either encodes a biologically relevant compound to LDL-cholesterol, or is a proxy for an existing or proposed LDL-cholesterol lowering drug. Genetic associations with LDL-cholesterol were obtained from the Global Lipids Genetics Consortium’s 2013 data release [17], and associations with CAD risk from CARDIoGRAMplusC4D’s 2015 data release [18]. These associations are displayed graphically in Figure 1 (left panel).

**Figure 1:**
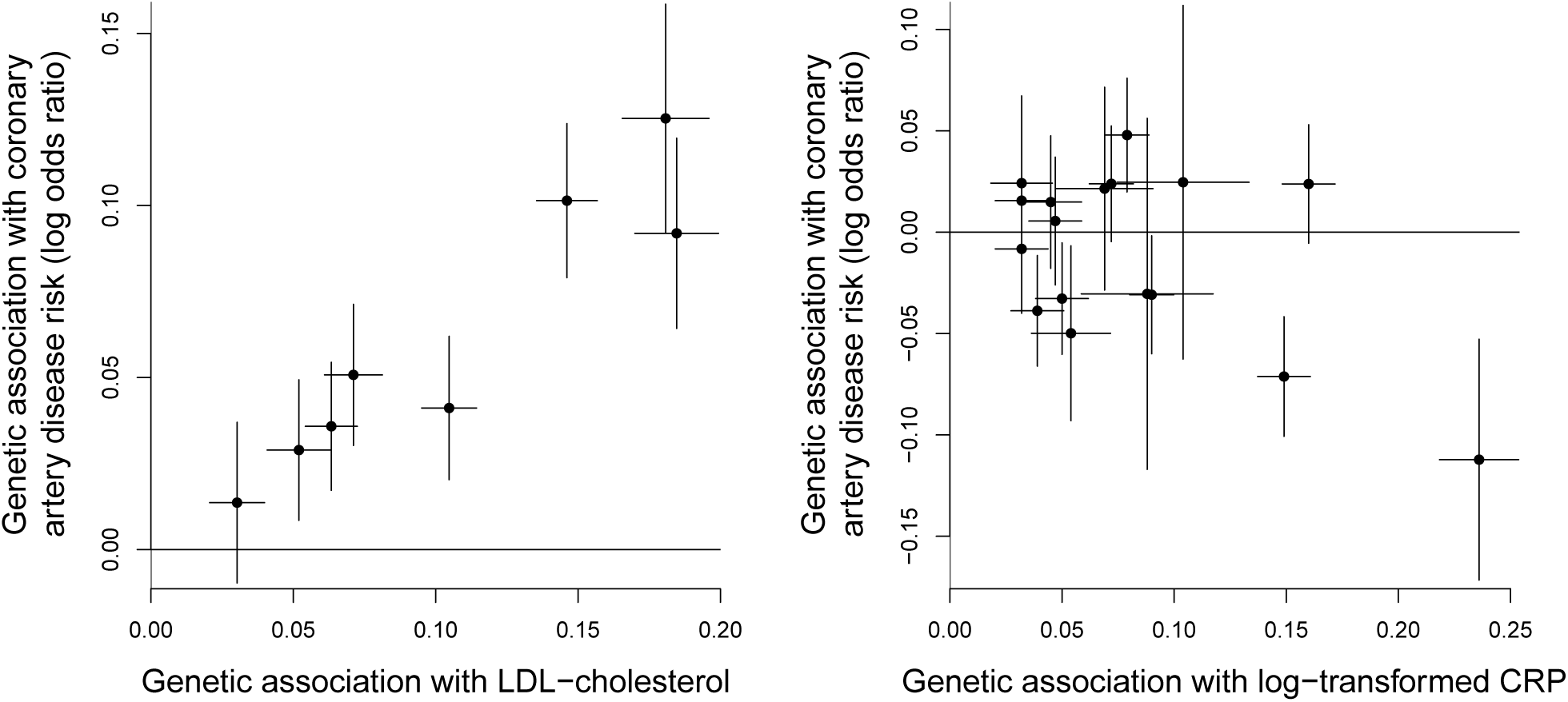
Genetic associations with risk factor and outcome (lines are 95% confidence intervals) for: (left) 8 genetic variants having biological links to LDL-cholesterol; (right) 17 genetic variants associated with C-reactive protein (CRP) at a genome-wide level of significance.

### C-reactive protein and CAD risk

We also consider the causal relationship between C-reactive protein (CRP) and CAD risk based on 17 genetic variants previously demonstrated to be associated with CRP at a genome-wide level of statistical significance [19]. The biological rationale for this analysis is not to evaluate the causal role of CRP, as several of these genetic variants are not specifically associated with CRP and hence are not valid instruments. The causal role of CRP can be evaluated in a Mendelian randomization analysis using genetic variants in the *CRP* gene region, the region that encodes CRP [20]. Rather, the biological rationale for this analysis considers CRP as a proxy measure for inflammation more generally, and investigates whether there are any consistent causal relationships between inflammation and CAD risk. Genetic associations with CRP are obtained from Dehghan et al. [19], and associations with CAD risk from the CARDIoGRAM consortium [21]. These associations are displayed graphically in Figure 1 (right panel).

### Results

Results for both examples are presented in Table 3. Estimates represent log odds ratios for CAD per 1 mmol/L increase in LDL-cholesterol, or per unit increase in log-transformed CRP. For the MBE method, we present estimates for a range of values of the bandwidth in the kernel-density estimator representing the suggested bandwidth from the modified Silverman rule (*ϕ* = 1), half the suggested bandwidth (*ϕ* = 0.5), and one-quarter of the suggested bandwidth (*ϕ* = 0.25), as well as for simple and weighted versions of the method.

**Table 3.** Estimates (standard errors, SE) and 95% confidence intervals (CI) from a variety of methods for applied examples. Abbreviation: MBE = mode-based estimate of Hartwig et al. [9].

| Risk factor: | LDL-cholesterol |  | C-reactive protein |  |
| --- | --- | --- | --- | --- |
| Method | Estimate (SE) | 95% CI | Estimate (SE) | 95% CI |
| Inverse-variance weighted | 0.585 (0.044) | 0.499, 0.671 | -0.135 (0.102) | -0.334, 0.065 |
| MR-Egger | 0.611 (0.100) | 0.415, 0.807 | -0.223 (0.198) | -0.611, 0.165 |
| Simple median | 0.561 (0.067) | 0.429, 0.693 | 0.118 (0.155) | -0.187, 0.422 |
| Weighted median | 0.585 (0.057) | 0.473, 0.697 | -0.303 (0.108) | -0.515, -0.092 |
| Simple MBE ($\phi = 1$) | 0.522 (0.105) | 0.316, 0.727 | 0.295 (0.372) | -0.433, 1.023 |
| Simple MBE ($\phi = 0.5$) | 0.700 (0.136) | 0.434, 0.966 | 0.285 (0.502) | -0.698, 1.269 |
| Simple MBE ($\phi = 0.25$) | 0.699 (0.147) | 0.411, 0.987 | 0.306 (0.510) | -0.694, 1.305 |
| Weighted MBE ($\phi = 1$) | 0.686 (0.096) | 0.498, 0.875 | -0.407 (0.152) | -0.705, -0.108 |
| Weighted MBE ($\phi = 0.5$) | 0.697 (0.140) | 0.423, 0.971 | -0.458 (0.112) | -0.678, -0.238 |
| Weighted MBE ($\phi = 0.25$) | 0.696 (0.140) | 0.421, 0.970 | -0.472 (0.218) | -0.898, -0.045 |
| Heterogeneity-penalized model averaging <sup>a</sup> | 0.598 | 0.475, 0.718 | -0.441 | -0.602, -0.257 and 0.038, 0.352 <sup>b</sup> |

<sup>a</sup>The heterogeneity-penalized model averaging method does not estimate a standard error. For the risk factor LDL-cholesterol, and assuming normality, the standard error would be 0.062.
<sup>b</sup>The confidence interval in this case is the union of two disjoint ranges.

In the first example, all of the methods suggest a positive causal effect. In the model averaging method, the weight of the estimate including all 8 variants is 12.1%, and estimates with 7 or more variants comprise 42.1% of the total weight (compared with 0.4% and 3.6% of the weight with no heterogeneity penalization – equal weights). The width of the confidence interval from the model averaging method is similar to that from the weighted median method, and narrower than that from all other methods except for the standard IVW method. Confidence intervals from the MBE method are considerably wider than those from other methods, and vary in size by up to 40% for the different choices of bandwidth considered here. In the second example, the methods give varied estimates. In particular, the simple MBE method gives a positive estimate, whereas the weighted MBE method gives a negative estimate with a confidence interval that excludes zero. In contrast, the model averaging method gives a negative estimate, but a confidence interval that includes both negative and positive values, although excludes zero – it includes two disjoint ranges of values. Again, the precision of the MBE estimates varied for different choices of bandwidth, in the most extreme comparison by almost a factor of two.

Figure 2 shows the mixture distributions of the IVW estimates based on all subsets of genetics variants using both equal weights (dashed line) and heterogeneity-penalized weights (solid line) weights from the model averaging method. For the first example, the equally and penalized weighted distributions are similar, as the IVW estimates based on all subsets of variants are similar. For the second example, the heterogeneity-penalized distribution differs substantially from distribution using equal weights and is bimodal, indicating that there are groups of variants having similar weight of evidence supporting both a positive and a negative causal effect, and suggesting that there are causal mechanisms linked with inflammation that have both protective and harmful effects on CAD risk. This explains the composite confidence interval including both positive and negative values. Only the model averaging method is able to capture this feature of the data.

**Figure 2:**
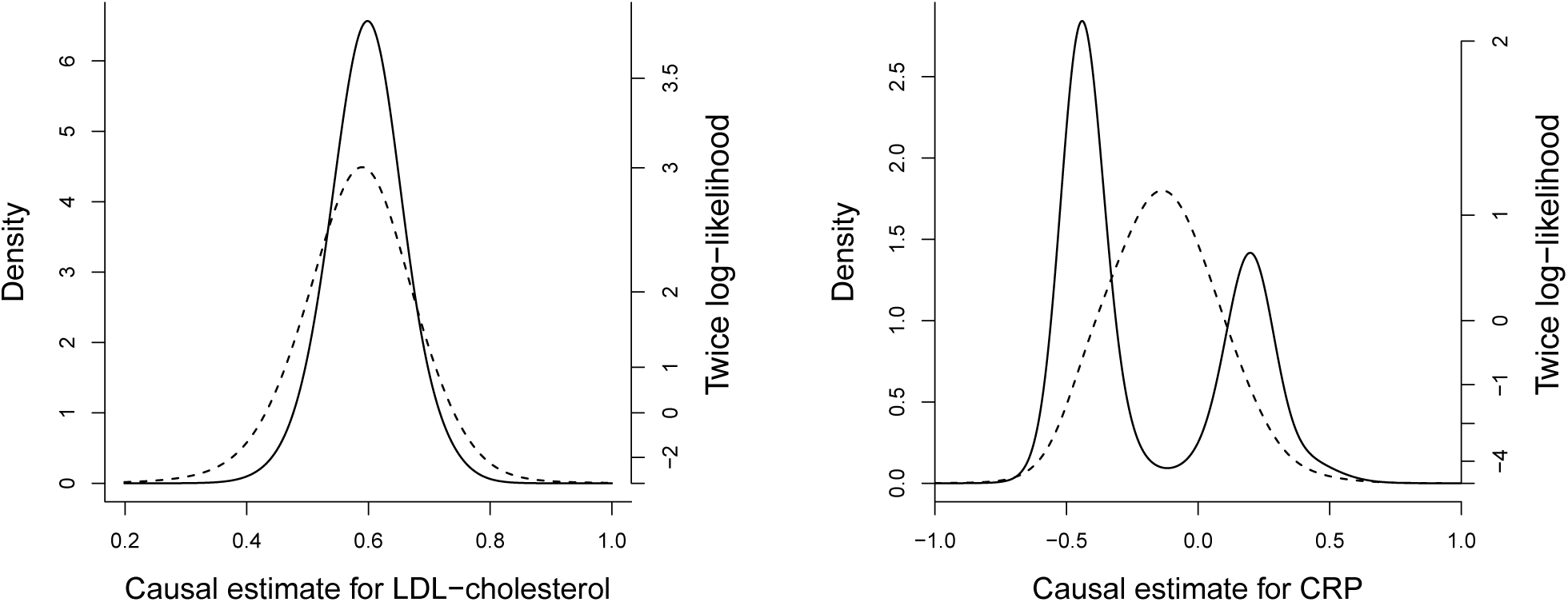
Mixture distributions of IVW estimates using equal (dashed line) and penalized (solid line) weights from model averaging method for: (left) LDL-cholesterol; (right) C-reactive protein (CRP). The right-hand axis is twice the log-likelihood – the 95% confidence interval contains all points within a vertical distance of 3.84 units on this scale (3.84 is the 95th percentile of a chi-squared distribution on 1 degree of freedom).

## Discussion

The aim of this manuscript was to develop a mode-based estimation method that provides a consistent estimate of the causal effect under the assumption that a plurality of the genetic variants are valid instruments. In comparison with the MBE method proposed by Hartwig et al., we believe that our method has several technical advantages: 1) it does not rely on the specification of a bandwidth parameter; 2) it makes inferences that do not rely on resampling methods; 3) it makes no asymptotic assumption about the distribution of the causal estimate for making inferences, in particular allowing confidence intervals to be asymmetric and to span multiple ranges; 4) it is asymptotically efficient, and should be efficient in finite samples, as the method seeks to upweight the IVW estimate based on the largest number of variants with homogeneous ratio estimates. One particular concern with the MBE method is that the precision of the estimate is highly variable depending on the choice of bandwidth parameter. There would be a great temptation as an applied researcher to perform the method for a variety of values of the bandwidth parameter, and choose the bandwidth parameter corresponding to the most desirable estimate.

The proposed heterogeneity-penalized model averaging method also outperformed Hartwig’s method in the simulation study, and in the applied examples. No sizeable inflation in Type 1 error rates was observed across the simulation scenarios when 2 or 3 of the 10 genetic variants were invalid, and bias and Type 1 error rates were generally either better or no worse than for other robust methods. The method was also at least as efficient as other robust methods when all variants were valid instruments, and had reasonable power to detect a causal effect throughout.

One deficiency of the proposed method is computational time. While the method was substantially quicker than that of Hartwig et al. with 10 genetic variants, the run-time of our method doubles with each additional variant. In the applied example with 17 genetic variants, 2^17^ − 1 = 131 071 weights were calculated. The method calculated weights in 0.7 seconds on a single 2.60 GHz central processing unit (CPU). The grid search algorithm took a 34 seconds. However, with 30 genetic variants, over 1 billion weights would need to be calculated. Reducing the computational burden may be possible – for example, models including genetic variants with highly discrepant ratio estimates would receive low weights and could be dropped with little loss of accuracy. However, solving this computational challenge in general is left as a problem for future work.

An extension of the method that could be valuable in applied practice is the use of prior information on particular variants. This can be achieved by multiplying the unnormalized weights *w*_*k*_ by a prior weighting *π*_0_(*k*) before normalizing. For example, if an investigator is particularly confident that a genetic variant is likely to be a valid instrument, then models containing this variant can be upweighted. Alternatively, prior weightings of models containing specific variants could be based on biological characteristics of the variants. For example, exonic and/or non-synonymous variants could be upweighted, or variants with functional information relating them to the risk factor. If these variants truly are more likely to be valid instruments, then this prior weighting would add to the robustness of the method. Additionally, a prior could be set to more strongly upweight less parsimonious models (that is, upweight models based on more genetic variants). This could add efficiency to the analysis, as models including more genetic variants will have more precise IVW estimates. Equal prior weights corresponds to a prior belief that 50% of genetic variants are valid instruments. If one instead believed that (say) 80% of genetic variants were valid instruments, then the prior for subset *k* could be set to *π*_0_(*k*) = 0.8^*K*^ *×* 0.2^*J-K*^ where *J* is the total number of genetic variants and *K* is the number of variants in subset *k*. The option to set this prior probability is included in the software code.

In conclusion, the heterogeneity-penalized model averaging procedure introduced in this paper will be a worthwhile contribution to the Mendelian randomization literature both in providing an additional robust method for causal estimation and testing the causal null hypothesis when some genetic variants may not be valid instruments, and for revealing features in the data such as the presence of multiple causal mechanisms.

## Acknowledgements

This work was supported by the UK Medical Research Council. Stephen Burgess and Verena Zuber are supported by Sir Henry Dale Fellowship jointly funded by the Wellcome Trust and the Royal Society (Grant Number 204623/Z/16/Z). Apostolos Gkatzionis is supported by a Medical Research Council Methodology Research Panel grant (Grant Number RG88311).

## Supplementary Material

### A.1 Software code

~~~
###### Summarized data set-up
bx # genetic associations with risk factor by # genetic associations with outcome
bxse # standard errors of genetic associations with risk factor byse # standard errors of genetic associations with outcome
######
bx =c(0.160, 0.236, 0.149, 0.09, 0.079, 0.072, 0.047, 0.05, 0.069,
  0.039, 0.088, 0.032, 0.104, 0.045, 0.054, 0.032, 0.032)
by =c(0.0237903, -0.1121942, -0.0711906, -0.030848, 0.0479207, 0.0238895,
  0.005528, -0.0327605, 0.0214852, -0.0387675, -0.0304042, -0.0082261,
  0.0246432, 0.0148795, -0.0498487, 0.0155667, 0.0242003)
bxse=c(0.006, 0.009, 0.006, 0.005, 0.005, 0.005, 0.006, 0.006, 0.011,
  0.006, 0.015, 0.006, 0.015, 0.007, 0.009, 0.006, 0.007)
byse=c(0.0149064, 0.0303084, 0.0150552, 0.0148339, 0.0143077, 0.0145478,
  0.0160765, 0.0140347, 0.0255237, 0.0139256, 0.0441698, 0.0162031,
  0.0444987, 0.016674, 0.0220043, 0.018098, 0.0219547)
  # example data (CRP-CAD associations)
  #
 ### Simple (but inefficient) code
  #
library(R.utils)
pen.weight < - function(theta, thetase, thetamean) {
 return (exp(-sum(log(thetase))-sum((theta-thetamean)^2/thetase^2/2))) }
  # this is the heterogeneity penalty weighting function
post=NULL; est=NULL; seest=NULL
  # these are the heterogeneity-penalized weights and means and standard deviations
  #  of the normal distributions in the weighted mixture distribution
for (i in 1:(2^length(bx)-1)) {
  inc=as.numeric(strsplit(intToBin(i),"")[[1]])
  inc=c(rep(0,length(bx)-length(inc)), inc)
  prior = ifelse(sum(inc)< 1.5, 0, 1)
  # prior is set to zero for all models with 0 or 1 variants,
  #  equal for all other subsets estinc = (by/bx)[which(inc==1)]
  seinc = abs((byse/bx)[which(inc==1)])
  meaninc = sum(estinc*seinc^-2)/sum(seinc^-2)
  weight = pen.weight(estinc, seinc, meaninc)
  post[i] = prior*weight
  est[i] = meaninc
  if (sum(inc) > 1) {
seest[i] = summary(lm(by[which(inc==1)]∼bx[which(inc==1)]-1,
     weights=byse[which(inc==1)]^-2))$coef[1,2]/
 min(summary(lm(by[which(inc==1)]∼bx[which(inc==1)]-1,
     weights=byse[which(inc==1)]^-2))$sigma, 1)
     }
 if (sum(inc) == 1) {
 seest[i] = byse[which(inc==1)]/bx[which(inc==1)] }
 }
post.norm = post/sum(post)
 # normalized heterogeneity-penalized weights
sumlik=NULL
point = seq(from=-1, to=1, by=0.001)
for (i in 1:length(point)) {
 lik = post.norm*dnorm(point[i], mean=est, sd=seest)
 sumlik[i] = sum(lik) }
  # calculates the likelihood at a range of values from -1 to +1
  # if the causal effect may be outside of this range,
  # then this range of values will need to be expanded
whichin = which(2*log(sumlik)> (2*max(log(sumlik))-qchisq(0.95, df=1)))
  # provides an index of estimate values in the 95% confidence interval estimate = -1.001+0.001*which.max(log(sumlik))
  # modal estimate
ifelse(sum(diff(whichin)!=1)==0, "Single range", "Multiple ranges")
  # returns "Single range" if the 95% CI is a single range of values
  # returns "Multiple ranges" otherwise
lowerCI = -1.001+0.001*whichin[1]
upperCI = -1.001+0.001*whichin[length(whichin)]
  # lower and upper confidence interval limits (assuming single range)
fullCI = -1.001+0.001*whichin
  # all estimate values in confidence interval
  # if the likelihood is calculated for a different range of values (not -1 to +1),
  # then this code will need to be altered
  #
  #
### Efficient (but harder to follow) code
  #
library(matrixStats);
library(iterpc);
library(Matrix);
library(stats);
library(optimbase);
  #
model.prior = function(model.size, N.obs, prob.valid.inst){
  pr = (prob.valid.inst^model.size)*(1-prob.valid.inst)^(N.obs-model.size)
  return(pr)
}
#
het.weight = function(prob.valid.inst, bx, by, byse){
  J = length(by);
  theta.est = by/bx;
  theta.se = abs(byse/bx);
  tmp.1 = by/byse;
  tmp.2 = bx/byse;
  theta.se.sq = theta.se^2;
  log.theta.se = log(theta.se);
  est = seest = vector("numeric", 2^J-1);
  het.weight = vector("numeric", 2^J-1);
  #
  count = 0;
 for(n in 1:J){
  perms = choose(J,n);
  inc = sparseMatrix(i=as.vector(t(replicate(n,1:perms))),
     j=as.vector(t(getall(iterpc(J,n,c(1:J))))),
     x=1, dims = c(perms,J));
   # sparse binary inclusion matrix
   # 1 denotes an instrument is included in the model
   # each row represents a particular model
   est.sum = inc%*%(theta.est/theta.se.sq);
   recip.var.ivw = inc%*%(1/theta.se.sq);
   est.ivw = est.sum/recip.var.ivw;
   est[(count+1):(count+perms)] = est.ivw;
if(n>1){
   tmp = t(replicate(J, as.vector(est.ivw)));
 if(n< J){
  psi.hat = sqrt((1/(n-1))*rowSums(t(t(inc)*(tmp.1^2 - 2*tmp*(tmp.1*tmp.2) + (tmp^2)*(tmp.2^2)))))
  }
 else{
  psi.hat = sqrt((1/(n-1))*sum(tmp.1^2 - 2*tmp*(tmp.1*tmp.2) + (tmp^2)*(tmp.2^2)));
  }
 psi.hat[which(psi.hat< 1)] = 1;
 seest[(count+1):(count+perms)] = psi.hat/sqrt(recip.var.ivw);
  }
else if(n==1){
 seest[(count+1):(count+perms)] = inc%*%theta.se;
       }
 #
if(n>1){
 het.exponent = rowSums(inc*t(t(t(t(inc)*theta.est) -
     as.vector(est.ivw))^2/theta.se.sq));
 het.weight[(count+1):(count+perms)] =
     exp(-(inc%*%(log.theta.se)+0.5*het.exponent))*
     model.prior(n,J,prob.valid.inst);
    }
 count = count+perms;
 } # ends for loop
 newlist = list(het.weight, est, seest);
 return(newlist)
 }
#
results = het.weight(0.5, bx, by, byse);
het.weight = results[[1]];
het.weight.norm = het.weight/sum(het.weight);
# normalized heterogeneity-penalized weights est = results[[2]];
seest = results[[3]];
  #
sumlik=NULL
grid.increment = 1e-3; grid.start = -1; grid.end = 1;
point = matrix(seq(grid.start, grid.end, grid.increment), ncol = 1);
  #
l = length(het.weight.norm);
sumlik = vapply(point,function(i){sum(het.weight.norm*dnorm(rep(i,l), est, seest))}, 1);
  # calculates the likelihood at a range of values from -1 to +1
  # if the causal effect may be outside of this range,
  # then this range of values will need to be expanded
whichin = which(2*log(sumlik)> (2*max(log(sumlik))-qchisq(0.95, df=1)));
  # provides an index of estimate values in the 95% confidence interval
estimate = -1.001+0.001*which.max(log(sumlik));
  # modal estimate
ifelse(sum(diff(whichin)!=1)==0, "Single range", "Multiple ranges");
  # returns "Single range" if the 95% CI is a single range of values
  # returns "Multiple ranges" otherwise lowerCI = -1.001+0.001*whichin[1];
upperCI = -1.001+0.001*whichin[length(whichin)];
# lower and upper confidence interval limits (assuming single range) fullCI = -1.001+0.001*whichin;
~~~

### A.2 Applied examples

#### LDL-cholesterol and CAD example

To assess the causal effect of LDL-cholesterol on CHD risk, we used 8 genetic variants in separate gene regions each of which has been specifically linked with LDL-cholesterol (each either encodes a biologically relevant compound to LDL-cholesterol, or is a proxy for an existing or proposed LDL-cholesterol lowering drug). These gene regions are: *HMGCR* (proxy for statin treatment), *PCSK9* (proxy for PCSK9 inhibition), *NPC1L1* (proxy for ezetimibe), *APOB* (encodes biologically relevant apolipoprotein B), *ABCG5/G8* (bile acid sequestrant), *SORT1* (antisense oligonucleotide RNA inhibitor targeting this pathway currently under development), *APOE* (encodes biologically relevant apolipoprotein E), and *LDLR* (encodes biologically relevant LDL receptor). The specific choice of variant in each gene region to include in the analysis was based on the lead variant from the 2010 analysis of the Global Lipids Genetic Consortium [Teslovich et al., Biological, clinical and population relevance of 95 loci for blood lipids. Nature 2010; 466:707–713].

Supplementary Table A1 provides information about these variants, including the beta-coefficients and standard errors for their associations per additional copy of the effect allele with LDL-cholesterol (mmol/L) and CAD risk (log odds ratios), together with the causal estimates based on each of these variants (log odds ratios for CAD per 1 mmol/L increase in LDL-cholesterol).

**Supplementary Table A1:** Details of genetic variants, beta-coefficients (standard errors, SE) for associations with low-density lipoprotein cholesterol (LDL-c, mmol/L) and with coronary artery disease (CAD) risk (log odds ratios) taken from CARDIoGRAM consortium, and causal effect estimates (log odds ratio per 1 mmol/L increase in LDL-cholesterol) for 8 genetic variants.

| rsid | Nearest gene | Effect allele | Association with LDL-c<br>Beta (SE) | Association with CAD risk<br>Beta (SE) | Causal estimate<br>Estimate (SE) |
| --- | --- | --- | --- | --- | --- |
| rs12916 | *HMGCR* | C | 0.063 (0.005) | 0.036 (0.009) | 0.566 (0.150) |
| rs2479409 | *PCSK9* | G | 0.052 (0.006) | 0.029 (0.010) | 0.556 (0.200) |
| rs2072183 | *NPC1L1* | C | 0.030 (0.005) | 0.014 (0.012) | 0.451 (0.394) |
| rs1367117 | *APOB* | A | 0.105 (0.005) | 0.041 (0.011) | 0.393 (0.101) |
| rs4299376 | *ABCG5/G8* | G | 0.071 (0.005) | 0.051 (0.010) | 0.714 (0.147) |
| rs629301 | *SORT1* | T | 0.146 (0.005) | 0.101 (0.011) | 0.694 (0.078) |
| rs4420638 | *APOE* | G | 0.185 (0.007) | 0.092 (0.014) | 0.498 (0.076) |
| rs6511720 | *LDLR* | G | 0.181 (0.008) | 0.125 (0.017) | 0.693 (0.094) |

#### CRP and CAD example

Supplementary Table A2 provides information about the 17 variants used in the example analysis of this paper for investigating causal relationships between inflammation and coronary artery disease (CAD) risk, using C-reactive protein (CRP) as a measure of inflammation. All variants were previously demonstrated to be associated with CRP levels at a genome-wide level of significance by Dehghan et al. [Meta-analysis of genome-wide association studies in > 80 000 subjects identifies multiple loci for C-reactive protein levels. Circulation 2011; 123(7):731–738] Details of these variants are given, including the beta-coefficients and standard errors for their associations with CRP (log-transformed) and CAD risk (log odds ratios), together with the causal estimates based on each of these variants (log odds ratios for CAD per unit increase in log-transformed CRP).

**Supplementary Table A2:** Details of genetic variants, beta-coefficients (standard errors, SE) for associations with C-reactive protein (CRP, log-transformed) and with coronary artery disease (CAD) risk, and causal effect estimates (log odds ratios for CAD per unit increase in log-transformed CRP) for 17 genome-wide significant variants.

| rsid | Nearest gene | Effect allele | Association with CRP<br>Beta (SE) | Association with CAD risk<br>Beta (SE) | Causal estimate<br>Estimate (SE) |
| --- | --- | --- | --- | --- | --- |
| rs2794520 | *CRP* | C | 0.160 (0.006) | 0.024 (0.015) | 0.149 (0.093) |
| rs4420638 | *APOC1* | A | 0.236 (0.009) | -0.112 (0.030) | -0.475 (0.128) |
| rs1183910 | *HNF1A* | G | 0.149 (0.006) | -0.071 (0.015) | -0.478 (0.101) |
| rs4420065 | *LEPR* | C | 0.090 (0.005) | -0.031 (0.015) | -0.343 (0.165) |
| rs4129267 | *IL6R* | C | 0.079 (0.005) | 0.048 (0.014) | 0.607 (0.181) |
| rs1260326 | *GCKR* | T | 0.072 (0.005) | 0.024 (0.015) | 0.332 (0.202) |
| rs12239046 | *NLRP3* | C | 0.047 (0.006) | 0.006 (0.016) | 0.118 (0.342) |
| rs6734238 | *IL1F10* | G | 0.050 (0.006) | -0.033 (0.014) | -0.655 (0.281) |
| rs9987289 | *PPP1R3B* | A | 0.069 (0.011) | 0.021 (0.026) | 0.311 (0.370) |
| rs10745954 | *ASCL1* | A | 0.039 (0.006) | -0.039 (0.014) | -0.994 (0.357) |
| rs1800961 | *HNF4A* | C | 0.088 (0.015) | -0.030 (0.044) | -0.346 (0.502) |
| rs340029 | *RORA* | T | 0.032 (0.006) | -0.008 (0.016) | -0.257 (0.506) |
| rs10521222 | *SALL1* | C | 0.104 (0.015) | 0.025 (0.044) | 0.237 (0.428) |
| rs12037222 | *PABPC4* | A | 0.045 (0.007) | 0.015 (0.017) | 0.331 (0.371) |
| rs13233571 | *BCL7B* | C | 0.054 (0.009) | -0.050 (0.022) | -0.923 (0.407) |
| rs2836878 | *PSMG1* | G | 0.032 (0.006) | 0.016 (0.018) | 0.486 (0.566) |
| rs4903031 | *RGS6* | G | 0.032 (0.007) | 0.024 (0.022) | 0.756 (0.686) |

